# Dissecting genetic and sex-specific host heterogeneity in pathogen transmission potential

**DOI:** 10.1101/733915

**Authors:** Jonathon A. Siva-Jothy, Pedro F. Vale

## Abstract

Heterogeneity in disease transmission is widespread and, when not accounted for, can produce unpredictable outbreaks of infectious disease. Despite this, precisely how different sources of variation in host traits drive heterogeneity in disease transmission is poorly understood. Here we dissected the sources of variation in pathogen transmission using *Drosophila melanogaster* and *Drosophila* C Virus as a host-pathogen model system. We found that infected lifespan, viral growth, virus shedding, and viral load at death were all significantly influenced by fly genetic background, sex and female mating status. To understand how variation in each of these traits may generate heterogeneity in disease transmission, we estimated individual transmission potential by integrating data on virus shedding and lifespan alongside previously collected data on social aggregation. We found that ∼15% of between-individual heterogeneity in disease transmission was explained by a significant interaction between genetic and sex-specific variation. We also characterised the amount of variation in viral load, virus shedding, and lifespan following infection that could be explained by genetic background and sex. Amongst the determinants of individual variation in disease transmission these sources of host variation play roles of varying importance, with genetic background generally playing the largest role. Our results highlight the importance of characterising sources of variation in multiple host traits when studying disease transmission at the individual-level.

## Introduction

Individual host heterogeneity in disease spread is commonly observed across a wide range of infectious diseases (Woolhouse et al., 1997; Lloyd-Smith et al., 2005; Paull et al., 2011). Such heterogeneity is so common that it has been generalised into the ‘20-80 rule’ because of the frequent observation that 20% of hosts contribute to roughly 80% of transmission (Shaw & Dobson, 1995; Wilson et al., 2002; Woolhouse et al., 1997). More extreme forms of heterogeneity can result in very rare ‘superspreading’ individuals capable of causing large outbreaks of infectious disease in human and animal populations (Brooks-Pollock, Roberts, & Keeling, 2014; Lloyd-Smith et al., 2005). A superspreader of particular infamy is Mary Mallon who became known as ‘Typhoid Mary’ by infecting over 50 people with *Salmonella typhi* while working as a cook in New York during the early 20^th^ century (Marineli, Tsoucalas, Karamanou, & Androutsos, 2013). More recently, the 2003 outbreaks of SARS in Singapore and Hong Kong were greatly accelerated by a few superspreading individuals who caused over 70% of all SARS transmission (Li et al., 2004).

Outbreaks of infectious disease are often difficult to predict, especially when the effect of superspreaders are not accounted for by traditional assessments of outbreak risk. A widely used metric for the rate of pathogen spread is the basic reproductive number, *R*_0_, which estimates the average number of expected secondary infections caused by a single infected individual in a completely susceptible population. By focussing on the population average, *R*_0_ conceals outliers with a potentially higher propensity to spread disease (Lloyd-Smith et al., 2005; Paull et al., 2012; VanderWaal & Ezenwa, 2016). A clearer understanding of what drives heterogeneity in disease transmission requires a framework capable of accounting for such between-individual variation, which could enable more efficient control strategies that specifically target and treat high-risk individuals (Lloyd-Smith et al., 2005). The importance of predicting high-risk individuals before outbreaks occur has pushed understanding the causes of heterogeneity in disease transmission to the forefront of epidemiology and disease ecology research (Gervasi, Civitello, Kilvitis, & Martin, 2015; Paull et al., 2012; Stein, 2011; VanderWaal & Ezenwa, 2016).

Despite being commonplace, the underlying causes of heterogeneity in pathogen transmission remain elusive. Individual variation in host contact networks may be an important factor: it was Typhoid Mary’s position as a cook which exposed her to so many susceptible individuals. However, what enabled Typhoid Mary to stay in this role was her status as an asymptomatic carrier of the infection, which led to her release from quarantine on several occasions (Marineli et al., 2013). Similarly, the absence of symptoms in a number of SARS superspreaders delayed their admission to hospital and allowed them to continue spreading the virus (Centers for Disease Control and Prevention (CDC), 2003). These examples help underline that achieving a detailed understanding of the sources of heterogeneity in pathogen transmission is challenging because it results from complex interactions between multiple host behavioural, physiological, and immune traits. By dissecting the underlying genetic and sex-specific sources of variation in these traits we can assess how they influence three key components of pathogen transmission: contact rate between infected and susceptible individuals, the likelihood that contact will result in infection, and the duration of infection (VanderWaal & Ezenwa, 2016).

Infected-susceptible host contact rate is predominantly determined by host behaviours affecting locomotion and aggregation. Contact rates are also affected by population density (Keeling & Rohani, 2007), social group size (Patterson & Ruckstuhl, 2013), and behavioural syndromes (Keiser, Pinter-Wollman, et al., 2016). Social networks often exhibit extreme heterogeneity in the wild (Godfrey, 2013; Rushmore et al., 2013) and factors such as host genotype, sex condition, age and personality have been demonstrated to affect social aggregation in lab systems (de Bono & Bargmann, 1998; Keiser, Howell, et al., 2016; Saltz, 2011; Siva-Jothy & Vale, 2019). Once individuals acquire an infection, their ability to clear and shed pathogens is chiefly determined by physiological and immune mechanisms. Variation in these mechanisms chiefly influence the likelihood of pathogen transmission and the duration of infection (Grassly & Fraser, 2008; VanderWaal & Ezenwa, 2016). Many genetic and environmental sources of variation in physiological immunity have been described (Bou Sleiman et al., 2015; Lazzaro Brian P & Little Tom J, 2009; Ponton et al., 2013) including coinfection (Budischak et al., 2015; Lass Sandra et al., 2013), nutrition (Cornet, Bichet, Larcombe, Faivre, & Sorci, 2014; Vale, Choisy, & Little, 2013), and stress (Beldomenico & Begon, 2010; Capitanio et al., 2008). It is relevant to note that most studies have addressed the effects of behavioural, physiological and immune traits on transmission in isolation of one another. However, there is increasing evidence that transmission heterogeneity is often explained by coupled heterogeneities in these traits and how they may covary (Bolzoni, Real, & Leo, 2007; Farrington, Whitaker, Unkel, & Pebody, 2013; White, Forester, & Craft, 2018). To fully understand the sources of heterogeneity in pathogen transmission, it is therefore essential to measure multiple behavioural, physiological, and immune traits in hosts.

In the present work we aimed to test how common sources of variation between individuals (genetic background, sex and mating status) contribute to individual heterogeneity in pathogen transmission potential. The fruit fly, *Drosophila melanogaster*, is a powerful and genetically tractable model of infection, immunity and behaviour (Apidianakis & Rahme, 2009; Sokolowski, 2001). This makes it an ideal model system to investigate heterogeneity in pathogen transmission in the highly controlled conditions of a laboratory. We infected males and females from a range of naturally derived genotypes with *Drosophila* C Virus (DCV), a horizontally transmitted fly pathogen that causes behavioural, physiological and metabolic pathologies (Arnold, Johnson, & White, 2013; Chtarbanova et al., 2014; Gupta, Stewart, Rund, Monteith, & Vale, 2017; Vale & Jardine, 2015). We then quantified host traits and infection outcomes that directly impact pathogen transmission: (1) the infected lifespan, (2) the internal viral load, (3) how much virus was shed, and (4) the viral load at death (VLAD). Finally, we integrated these measurements alongside previously described data on variation in social aggregation (Siva-Jothy & Vale, 2019) into a composite metric of individual transmission potential, *V* (Lloyd-Smith et al., 2005; VanderWaal & Ezenwa, 2016). Estimations of individual transmission potential, *V*, allowed us to assess how genetic and sex-specific variation affects between-individual heterogeneity in pathogen transmission.

## Materials & Methods

### Flies & Rearing Conditions

Flies used in experiments were 3-5 days old and came from ten lines of the *Drosophila* Genetic Resource Panel (DGRP). These genetic backgrounds are five of the most resistant and susceptible to systemic Drosophila C Virus infection (Magwire et al., 2012). Virgin females were isolated from males within 7 hours of eclosion. Mated females and males were produced by rearing one female with one male for 24 hours. Mating was confirmed using oviposition within the following 24 hours and these egg’s subsequent development. Flies were reared in plastic vials on a standard diet of Lewis medium at 18±1°C with a 12 hour light:dark cycle. Stocks were tipped into new vials approximately every 14 days. One month before the experiments, flies were maintained at low density (∼10 flies per vial) for two generations at 25±1°C with a 12 hour light:dark cycle.

### Virus Culture & Infection

The *Drosophila* C Virus (DCV) isolate used in this experiment was originally isolated in Charolles, France and grown in Schneider *Drosophila* Line 2 (DL2) as previously described (Vale and Jardine, 2015b), diluted ten-fold (10^8^ infectious units per ml) in TRIS-HCl solution (pH=7.3), aliquoted and frozen at -70°C until required. To infect with DCV, flies were pricked in the pleural suture with a 0.15mm diameter pin, bent at 90° ∼0.5mm from the tip, dipped in DCV.

### Measuring Lifespan and Viral Load at Death

Lifespan and viral load at death were measured in the same fly. Following DCV infection, flies were isolated and reared in standard vials. Flies were then monitored every day until all individuals died, whereupon they were removed from vials, fixed in 50μl of TRI-reagent and frozen at -70°C, to await DCV titre at death quantification. For twenty-seven of thirty treatment groups, the lifespan following infection and viral load at death was measured for n=17-20, three treatment groups consisted of n=7-15 flies (Table S1).

### Viral Growth and Shedding Measurement Setup

Due to destructive sampling, we measured the viral load and shedding of single flies at a single time point, either 1-, 2- or 3-days post-infection (DPI). Following DCV infection, single flies were placed into 1.5ml Eppendorf tubes with ∼50μl of Lewis medium in the bottom of the tube. To measure viral shedding, flies were transferred to tubes for 24 hours, immediately following 1 or 2 days after systemic infection. After living in these tubes for a further 24 hours, flies were removed and homogenised in 50μl of TRI-reagent, tubes were also washed out with 50μl of TRI-reagent by vortexing. These samples were then frozen at -70°C, to await DCV quantification by qPCR. For each combination of sex and genetic background over the three days vial load and virus shedding was measured, n=7-15 flies were measured (Table S2-S4).

### DCV RNA Extraction

RNA was extracted from viral load at death and viral shedding samples by Phenol-Chloroform extraction. Samples were thawed on ice for 30 minutes before being incubated at room temperature for 5 minutes to allow dissociation of nucleo-protein complex. Samples were then centrifuged at 12,000×g for 10 minutes at 4°C after which large debris was removed. For phase separation, samples were shaken vigorously for 15 seconds, 10μl of chloroform added, incubated at room temperature for a further 3 minutes before being centrifuged at 12,000×g for 15 minutes at 4°C. Following phase separation, the upper aqueous layer was removed from each sample and added to 25μl of isopropanol, tubes were then inverted twice, before being centrifuged at 12,000×g for 10 minutes at 4°C. Precipitated RNA was then washed by removing the supernatant, and re-dissolving the RNA pellet in 50μl of 75% ethanol before being centrifuged at 7,500×g for 5 minutes at 4°C. RNA suspension was achieved by removing 40μl of the ethanol supernatant, allowing the rest to dry by evaporation and dissolving the remaining RNA pellet in 20μl of RNase-free water. We extracted RNA from flies after 1, 2 or 3 days of infection using a semi-automatic MagMAX Express Particle Processor using the MagMAX-96 total RNA isolation kit manufacturer’s protocol (Life Technologies, 2011) with the elution step extended to 18 minutes. RNA samples were stored at -70°C to await reverse transcription.

### Reverse transcription & qPCR Protocol

Extracted RNA was reverse-transcribed with M-MLV reverse transcriptase and random hexamer primers, before being diluted 1:1 with nuclease free water. cDNA samples were stored at -20°C to await qPCR analysis. DCV titre was quantified by qPCR using Fast SYBR Green Master Mix in an Applied Biosystems StepOnePlus system. Samples were exposed to a PCR cycle of 95°C for 2 minutes followed by 40 cycles of: 95°C for 10 seconds followed by 60°C for 30 seconds. Forward and reverse primers used included 5’-AT rich flaps to improve fluorescence (DCV_Forward: 5’ AATAAATCATAAGCCACTGTGATTGATACAACAGAC 3’; DCV Reverse: 5’ AATAAATCATAAGAAGCACGATACTTCTTCCAAACC 3’). Across all plates, two technical replicates were carried out per sample. DCV titre was calculated by absolute quantification, using a standard curve created from a 10-fold serial dilution (1-10^-12^) of DCV cDNA. Our detection threshold was calculated for each plate using the point at which two samples on our standard curve gave the same Ct value. The point of redundancy in a standard curve was taken to be equivalent to 0 viral particles. Due to our detection protocol measuring viral copies of RNA, we cannot comment on the viability of any detected virus. We transformed our measurements of viral RNA transformed in order for them to represent the amount of virus growing inside a whole fly rather than the amount in the qPCR well sample. To account for dilution between RNA extraction and qPCR we transformed DCV RNA samples by a factor of 3125, to represent the amount of DCV growing in, or shed by, flies. The mean qPCR efficiency was 116% with a standard error of ±2.9%

### Calculating Between-Individual Variation in Transmission Potential, *V*

We used measures of virus shedding, lifespan following infection, and social aggregation to predict individual transmission potential. We integrated these measures based on a simple framework that describes transmission potential as a function of contact rate between susceptible and infected individuals, the likelihood that such contact will result in infection, and the duration of the infectious period (VanderWaal & Ezenwa, 2016). Using previously analysed data on social aggregation (Siva-Jothy & Vale, 2019), we used nearest neighbour distance as a measure of contact rate. Flies that aggregated more closely to conspecifics, have a higher contact rate, and are therefore more likely to spread DCV. We also assume that transmission likelihood increases with virus shedding. We therefore take the amount of virus shed by flies as a proximate measure of the likelihood that contact will result in infection. Using these traits, individual transmission potential, *V*, was calculated as:

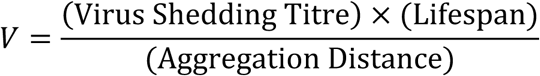

Aggregation distance, lifespan following infection and virus shedding were all measured in separate experiments. Therefore, to calculate V as a measure of individual transmission potential, we simulated theoretical individuals by bootstrapping trait values sampled from each of these three datasets. We simulated 60 individuals for each combination of sex and genetic background, assuming no specific covariance structure between traits, that is, all possible trait combinations were considered.

### Statistical Analysis

Across all experiments, generalised linear models (GLMs) were used to analyse continuous response variables and logistic regressions were used to analyse proportions. An effect of sex or mating was analysed in separate models comparing males or virgin females to the same dataset of mated females, respectively.

To analyse lifespan, two GLMs were constructed containing a three-way interaction genetic background, VLAD, and sex or mating (Table S6). The two GLMs for VLAD, contained either a two-way interaction between genetic background and sex or a two-way interaction between genetic background and mating (Table S6).

Due to zero-inflation, we used two models to sequentially analyse both viral load and virus shedding data. Viral load and virus shedding are broken down into qualitative (the proportion of non-zero values) and quantitative variation (differences between non-zero values). First, we conducted logistic regressions on all of the values in these datasets and analysed the proportion of values that were greater than zero. Logistic regressions analysing sex-differences in viral load included DPI (a 3-level factor: 1, 2 or 3 days) and an interaction between genetic background and sex (Table S6). For analysing the effect of mating in females on viral load, logistic regressions included DPI and an interaction between genetic background and mating (Table S6). Logistic regressions of virus shedding used a similar model that also included quantitative viral load as a predictor (Table S6). After these logistic regressions, zeroes were removed from all datasets to analyse the subset of positive-values. The GLMs used to analyse these subsets included the same predictors as their corresponding logistic regressions, for viral load: an interaction between genetic background and sex or mating, alongside DPI, with the inclusion of quantitative viral load for virus shedding (Table S6).

Due to zero-inflation *V* was also analysed with a logistic regression followed by a GLM. A logistic regression was used to analyse the proportion of *V* values that were greater than zero with a two-way interaction between sex and genetic background as predictors (Table S6). Zero-values of *V* were then removed from the dataset, and a GLM was used to analyse differences in the size of *V*, with an interaction between sex and genetic background included as a predictor (Table S6).

We calculated the amount of deviance and variance explained by predictors in logistic regressions and GLMs, respectively, by dividing the total deviance or variance explained by the model. Where appropriate, we corrected for multiple testing using Bonferroni correction. All statistical analyses and graphics produced in R 3.3.0 using the *ggplot2* (Wickham, 2016), *lme4* (Bates, Mächler, Bolker, & Walker, 2015) and *multcomp* (Hothorn, Bretz, & Westfall, 2008) packages.

## Results

### Lifespan Following Infection

Infected lifespan varied significantly between males and females and the extent of this variation differed between host genetic backgrounds (Figure 1a; Table 1). Genetic background explained the most variance of any predictor across models assessing mating (7%) and sex (10.9%). Interactions with sex and mating also explained a further 2.7% and 1.5%, respectively (Figure 5; Table 1). We found no evidence that mating affected the lifespan of females following DCV infection (Figure 1a; Table 1). Viral load at death (VLAD) was not affected by genetic background, sex or female mating status (Figure 1b; Table 2), and flies that died sooner following infection had greater VLAD than those that died later (Figure 1c; Table 1).

**Table 1.** Model outputs for the generalized linear modelling tests performed on lifespan following DCV infection. The VLAD acronym is used in place of ‘viral load at death’. Separate analyses were used to test the effect of sex and mating in females.

| Response Variable | Predictor | Df | F | %Variance Explained | p-value |
| --- | --- | --- | --- | --- | --- |
| Lifespan Following Infection | Sex | 1 | 2.00 | 0.6 | 0.16 |
|  | Genetic Background | 9 | 3.92 | 10.9 | <0.0001 |
|  | VLAD | 1 | 38.9 | 12.1 | <0.0001 |
|  | Sex*Genetic Background | 9 | 0.96 | 2.7 | 0.47 |
|  | Sex*VLAD | 1 | 5.46 | 1.7 | 0.02 |
|  | Genetic Background*VLAD | 9 | 0.63 | 1.8 | 0.77 |
|  | Sex*Genetic Background*VLAD | 9 | 2.67 | 7.4 | 0.005 |
|  | Mating | 1 | 2.74 | 0.9 | 0.099 |
|  | Genetic Background | 8 | 2.43 | 7.0 | 0.01 |
|  | VLAD | 1 | 32.3 | 10.2 | <0.0001 |
|  | Mating*Genetic Background | 8 | 0.54 | 1.5 | 0.84 |
|  | Mating*VLAD | 1 | 3.78 | 1.2 | 0.053 |
|  | Genetic Background*VLAD | 8 | 1.71 | 4.9 | 0.087 |
|  | Mating*Genetic Background*VLAD | 8 | 1.46 | 4.2 | 0.16 |

**Table 2.**
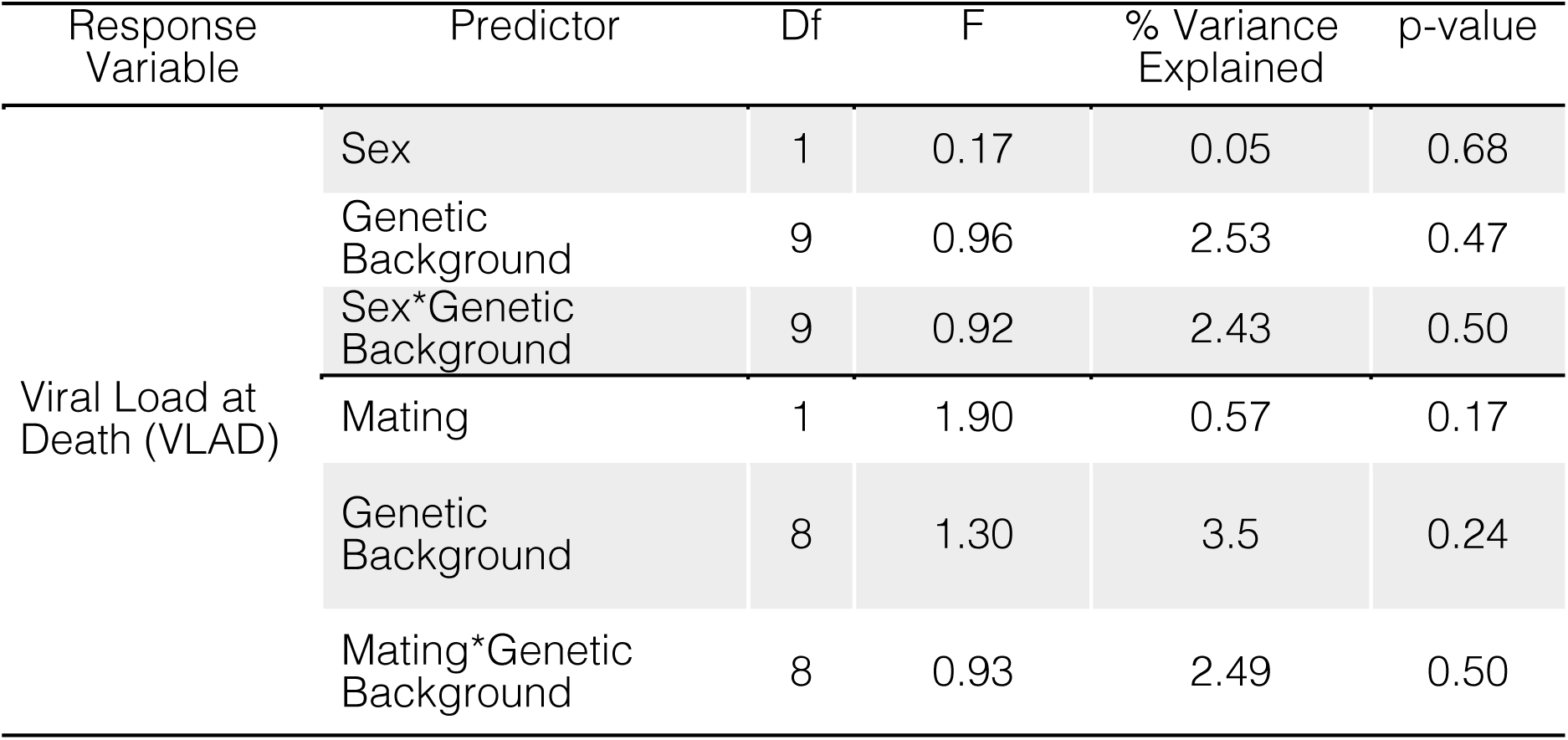
Model outputs for the generalized linear modelling tests performed on the viral load at death of flies infected with DCV. Separate analyses were used to test the effect of sex and mating in females.

**Figure 1.**
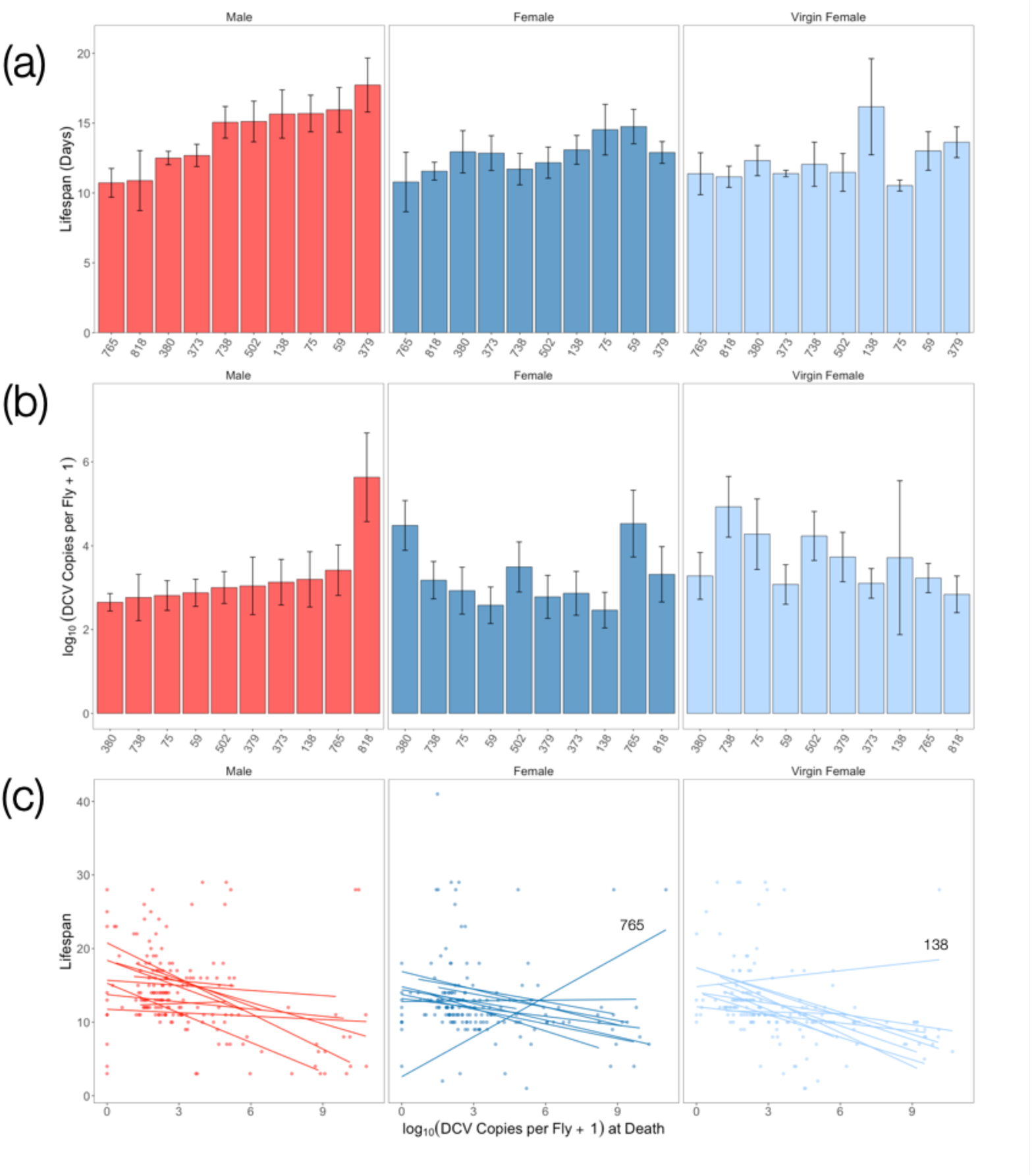
Mean±SE (a) lifespan in days following infection and (b) the viral load at death in males (red), mated females (blue), and virgin females (pale blue) of ten genetic backgrounds. The x-axis shows the line number form the DGRP panel and is in ascending order according to male flies. (c) the relationship between lifespan following infection and the viral load of flies at death. Each point is an individual male (red), mated female (blue), or virgin female (pale blue) fly. The nature of this relationship within each genetic background is represented by a line of best fit with outlier backgrounds labelled.

**Figure 2.**
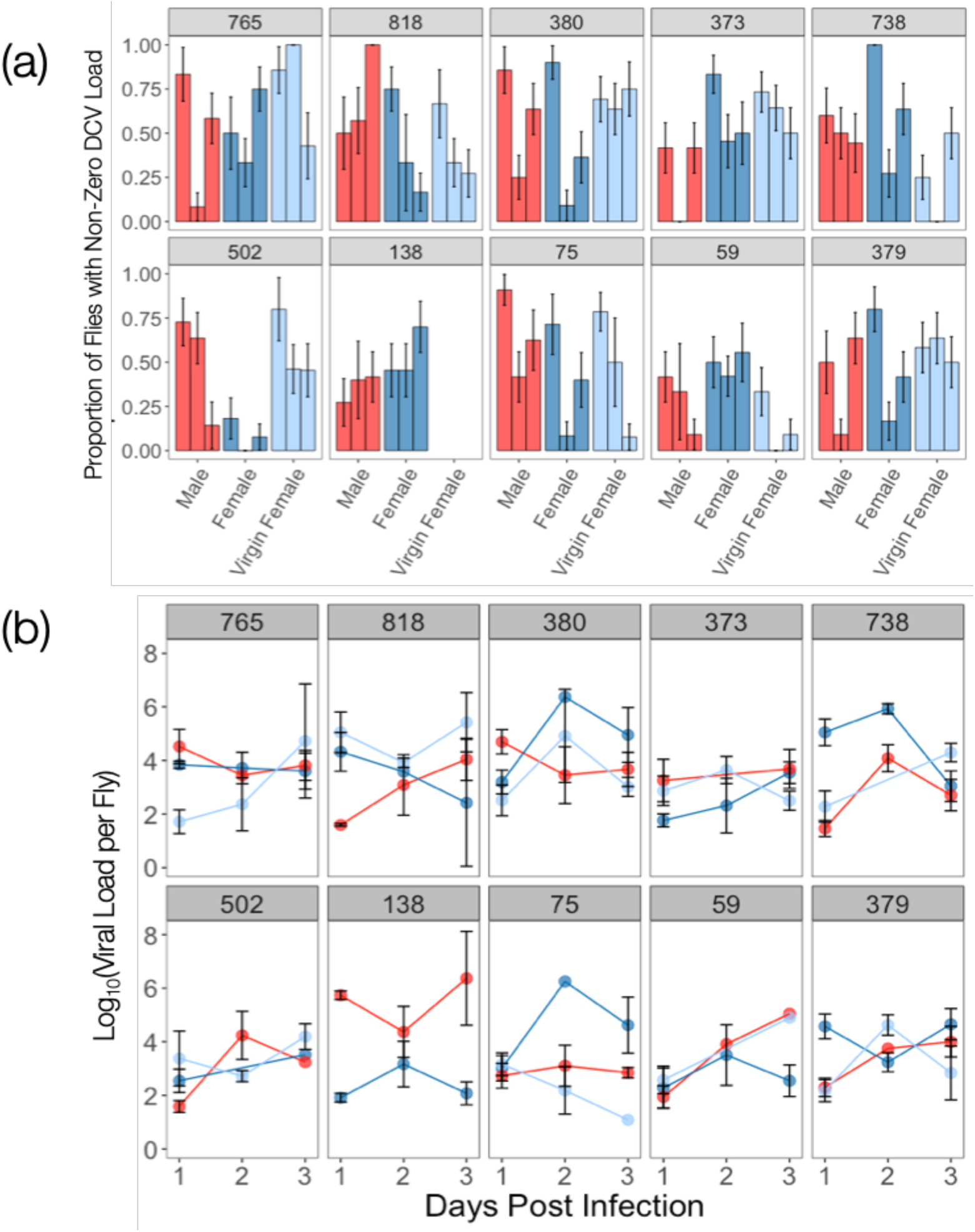
Mean±SE (a) proportion of flies with non-zero loads of DCV and the (b) viral titre of flies with non-zero DCV loads, over the first 3 days of infection. Across both panels, numbers in each pane denote the genetic background from the DGRP, while the colour of bars, points and lines represent sex and mating status. Males are shown in red, mated females in blue, and virgin females in pale blue. Bars of the same colour in each in pane in panel (a) represent (from left to right) days 1, 2 and 3 of infection.

**Figure 3.**
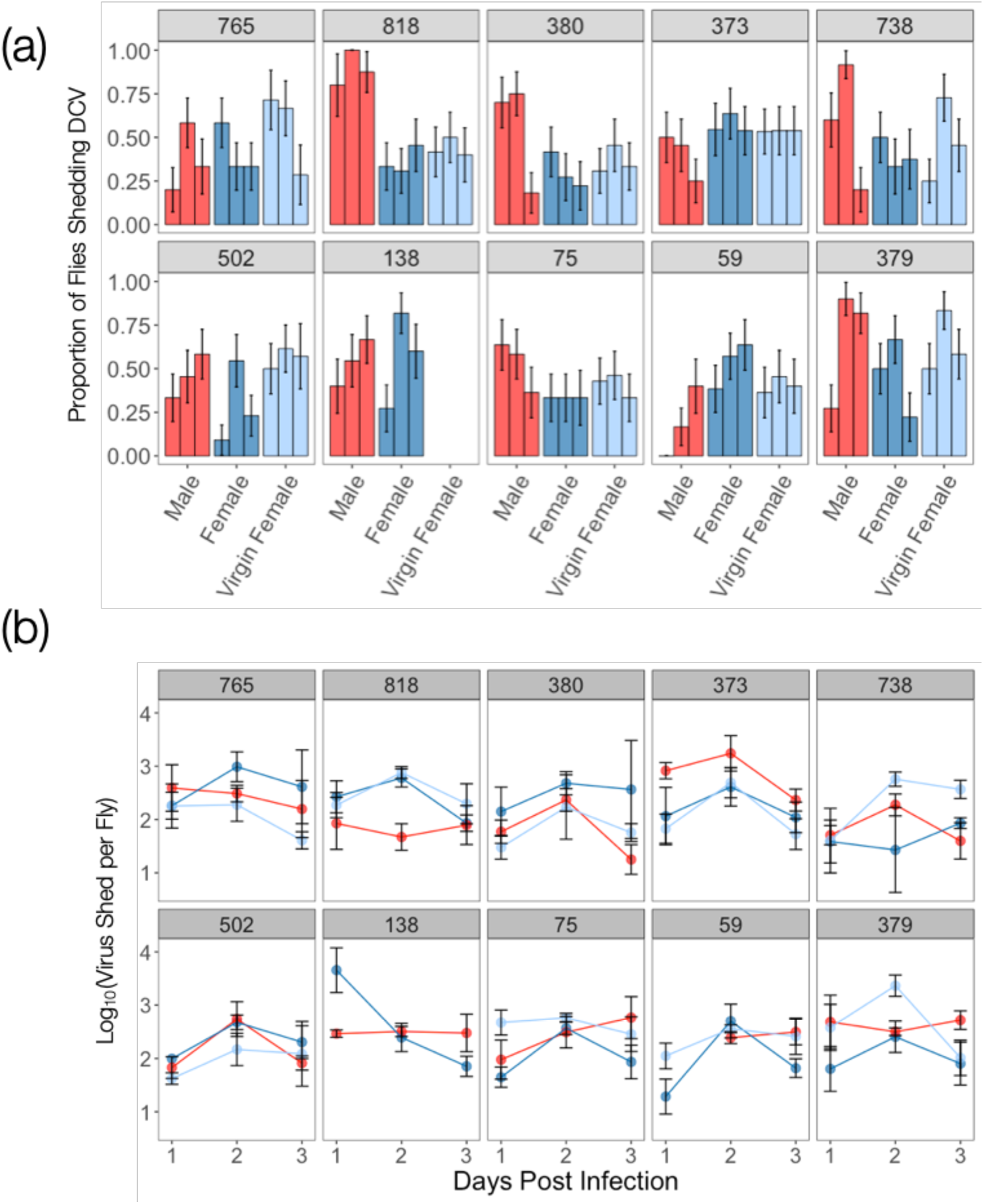
Mean±SE (a) proportion of flies shedding non-zero titres of DCV and the (b) titre of the non-zero virus shedding subset over the first 3 days of infection. Panels denote genetic background, while the colour of bars, points and lines represent sex and mating status. Males are shown in red, mated females in blue, and virgin females in pale blue. Bars of the same colour in each in pane in panel (a) represent (from left to right) days 1, 2 and 3 of infection.

**Figure 4.**
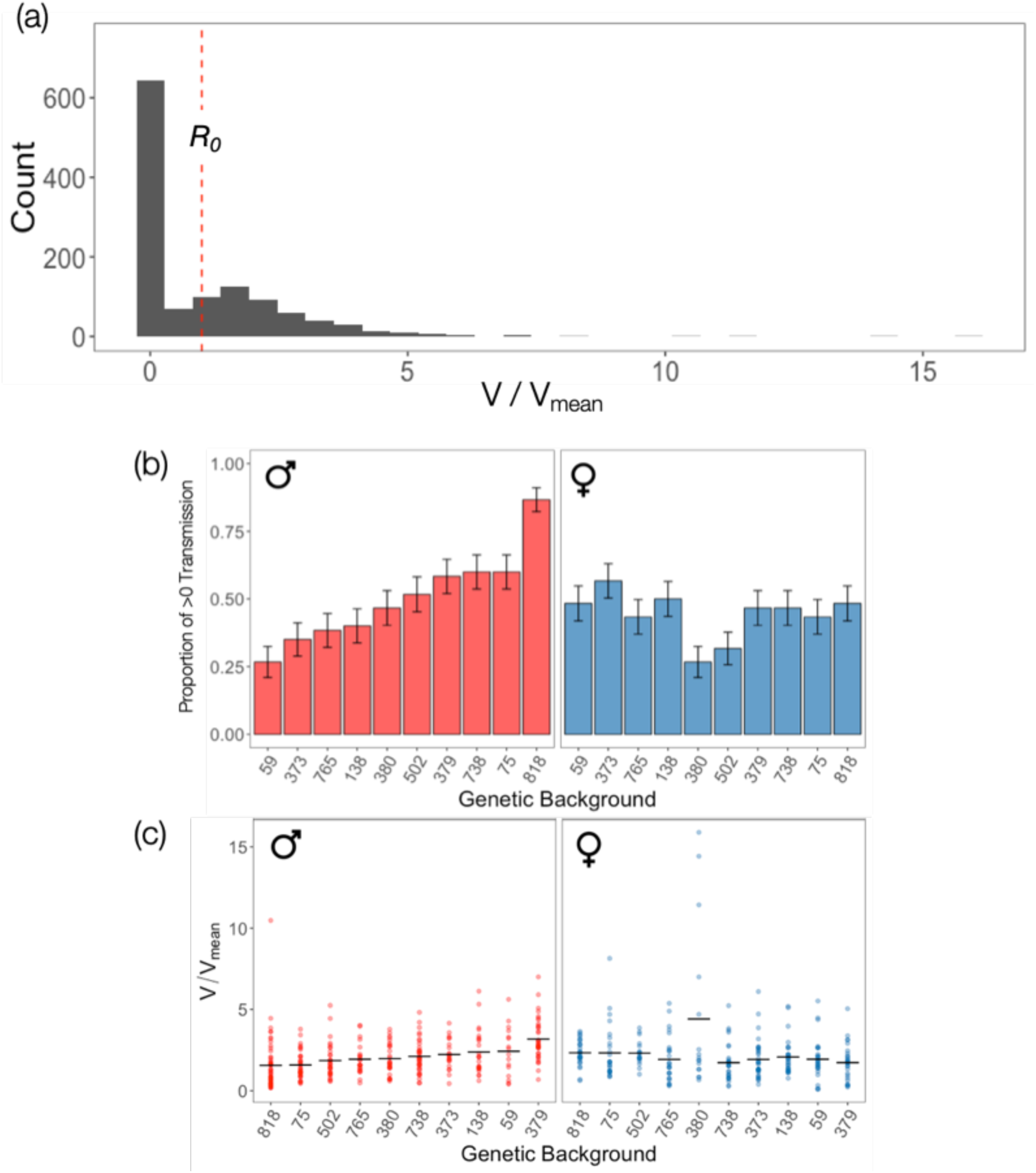
Bootstrap simulation results of transmission potential (*V*) (n=60): (a) the population-level distribution of *V* relative to the mean of the population. The red dashed line demarcates the average transmission potential of the population (similar to *R*_0_), a traditional metric used to describe a population’s outbreak risk. The mean±SE of (b) the proportion of flies with a non-zero transmission potential and (c) the transmission potential of flies with a non-zero transmission potential. In figure panels (b) and (c) sex is denoted by colour with males in red and females in blue. The x-axis of panels (b) and (c) is in ascending order of the male genetic backgrounds.

**Figure 5.**
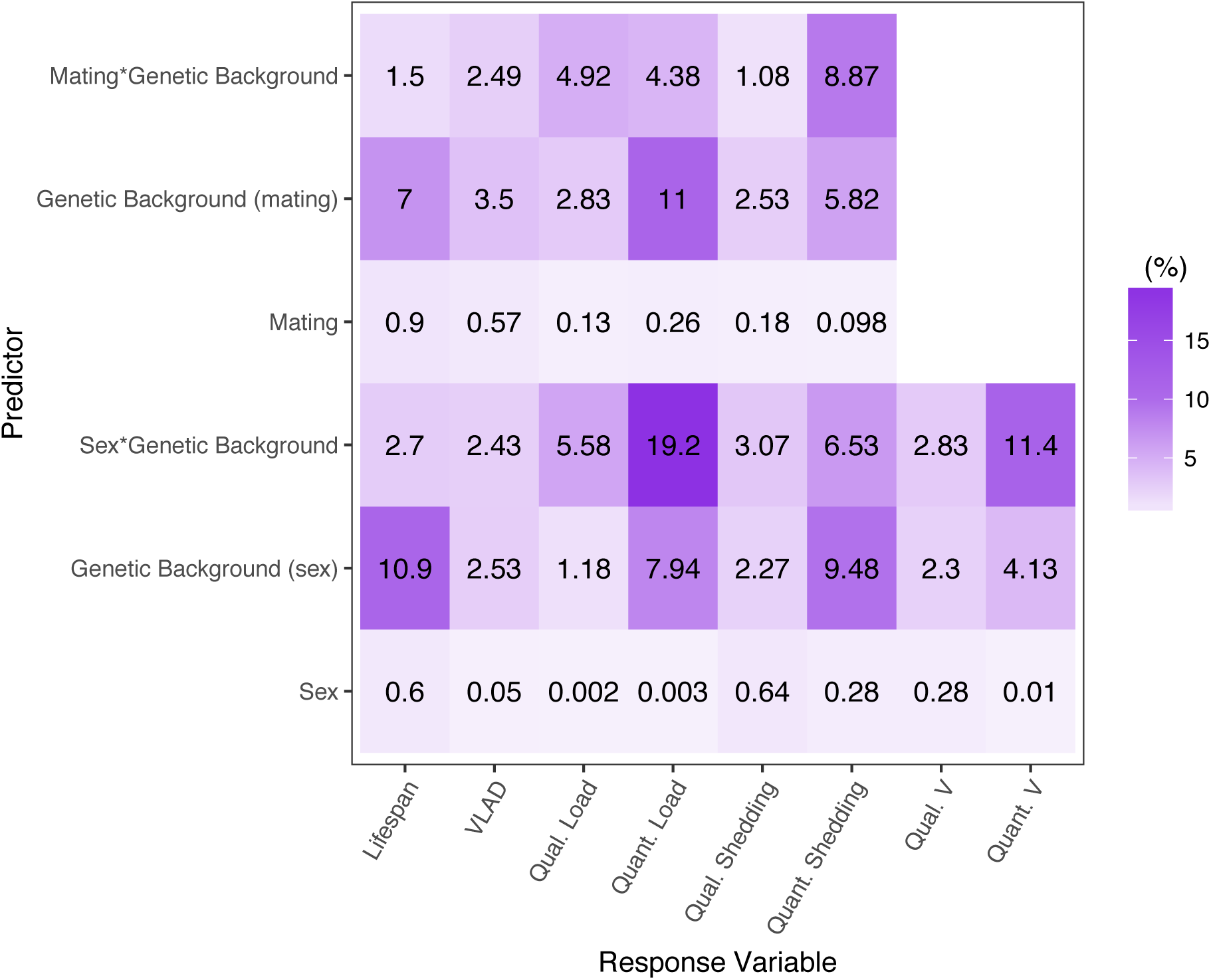
Summary of the percentage of variance or deviance explained by a subset of predictors in analyses of disease transmission potential and outcomes of infection.

### Viral Load

A substantial number of flies did not have detectable loads of DCV. These zero-values reflect qualitative variation and are likely caused by viral titres below the detection threshold of our qPCR and therefore reflect individuals with very low DCV loads, or no virus at all. We found extensive genetic variation in qualitative DCV load (Figure 2a; Table 3) which was affected by sex (Figure 2a; Table 3) and female mating status (Figure 2a; Table 3). Relatively little deviance was explained by sex (0.002%), mating (0.13%), or genetic background in models testing sex (1.18%) and mating (2.83%) effects. The predictors that explained the most deviance were the interactions between genetic background and sex (5.58%) or mating (4.92%) (Figure 5; Table 3). The size of non-zero DCV loads reflects quantitative variation and was affected by similar interactions between mating and sex with genetic background (Figure 2b; Table 4). While <1% of variance was explained by sex and mating, much more was explained by genetic background (7.94% and 11%) alongside its interactions with sex (19.2%) and mating (4.38%) (Figure 5; Table 4).

**Table 3.** Model outputs for the binomial logistic regression conducted on qualitative DCV loads (the proportion of non-zero DCV loads). The DPI acronym is used in place of ‘days post-infection’. Separate analyses were used to test the effect of sex and mating in females.

| Response Variable | Predictor | Df | $\chi^2$ | % Deviance Explained | $p$-value |
| --- | --- | --- | --- | --- | --- |
| Qualitative DCV Load | Sex | 1 | 0.019 | 0.002 | 0.89 |
|  | Genetic Background | 9 | 9.58 | 1.18 | 0.39 |
|  | DPI | 2 | 36.6 | 4.52 | <0.0001 |
|  | Sex*Genetic Background | 9 | 45.2 | 5.58 | <0.0001 |
|  | Mating | 1 | 1.01 | 0.13 | 0.31 |
|  | Genetic Background | 8 | 22.4 | 2.83 | 0.008 |
|  | DPI | 2 | 27.2 | 3.43 | <0.0001 |
|  | Mating*Genetic Background | 8 | 39.0 | 4.92 | <0.0001 |

**Table 4.** Model outputs for the GLM analysis conducted on quantitative DCV load (the titres of non-zero DCV loads). The DPI acronym is used in place of ‘days post-infection’. Separate analyses were used to test the effect of sex and mating in females.

| Response Variable | Predictor | DF | F | % Variance Explained | p-value |
| --- | --- | --- | --- | --- | --- |
| Quantitative DCV Load | Sex | 1 | 0.0062 | 0.003 | 0.94 |
|  | Genetic Background | 9 | 2.24 | 7.94 | 0.02 |
|  | DPI | 2 | 3.37 | 2.65 | 0.036 |
|  | Sex* Genetic Background | 9 | 5.41 | 19.2 | <0.0001 |
|  | Mating | 1 | 0.68 | 0.26 | 0.41 |
|  | Genetic Background | 8 | 3.18 | 11.0 | 0.0012 |
|  | DPI | 2 | 4.66 | 3.60 | 0.01 |
|  | Mating* Genetic Background | 8 | 1.42 | 4.38 | 0.19 |

The number of detectable DCV loads decreased following 1-day post-infection (pairwise comparison, p<0.0001) and remained lower than day 1 at day 3 (pairwise comparison, p=0.0016). There were significant changes in quantitative DCV load variation over the first three days of infection. Viral load peaked 2-days post-infection (pairwise comparison, p=0.0012), before decreasing to the same level as 1-day post-infection at 3-days post-infection (pairwise comparison, p=0.068).

### Virus Shedding

Similar to measures of viral load, we did not detect DCV in the shedding of a number of flies. Here, we interpret zeroes to be reflective of individuals that shed very low titres of DCV, or no virus at all. Qualitative variation in DCV shedding was significantly affected by genetic background, with sex modulating the extent of this difference (Figure 3a; Table 5). Sex however, explained <1% of the deviance, while genetic background and its interaction with sex explained 2.2% and 3.07% (Figure 5). Mating did not affect qualitative DCV shedding (Figure 3a; Table 5) and explained <1% of the deviance (Figure 5; Table 5). In flies where DCV was detected in shedding, quantitative DCV shedding was affected by genetic background and the extent of this variation was determined by female mating status, but not sex (Figure 3b; Table 6). The amount of variance explained by sex and in our models was <1%, in comparison with genetic background (9.48% and 5.82%) and its interactions with sex (8.87%) or mating (6.53%) (Figure 5; Table 6). Qualitative and quantitative DCV shedding peaked at day 2 (Figures 3a; Tables 5 & 6, pairwise comparisons, p<0.0001). Across all treatment groups, there was no significant relationship between viral load and shedding (Figure S1; Table 6).

**Table 5.** Model outputs for the GLM analysis conducted on qualitative DCV shedding (the proportion of sheddings with non-zero readings of DCV). The DPI acronym is used in place of ‘days post-infection’. Separate analyses were used to test the effect of sex and mating in females.

| Response Variable | Predictor | Df | $\chi^2$ | % Deviance Explained | p-value |
| --- | --- | --- | --- | --- | --- |
| Qualitative DCV Shedding | Sex | 1 | 4.93 | 0.64 | 0.026 |
|  | Genetic Background | 9 | 17.6 | 2.27 | 0.04 |
|  | Viral Load | 1 | 0.03 | 0.004 | 0.85 |
|  | DPI | 2 | 25.1 | 3.25 | <0.0001 |
|  | Sex*Genetic Background | 9 | 23.8 | 3.07 | 0.005 |
|  | Mating | 1 | 1.33 | 0.18 | 0.25 |
|  | Genetic Background | 8 | 19.0 | 2.53 | 0.025 |
|  | Viral Load | 1 | 1.10 | 0.15 | 0.29 |
|  | DPI | 2 | 7.66 | 1.02 | 0.022 |
|  | Mating*Genetic Background | 8 | 8.12 | 1.08 | 0.42 |

**Table 6.** Model outputs for the GLM analysis conducted on quantitative DCV shedding (the subset of shedding with non-zero readings of DCV). The DPI acronym is used in place of ‘days post-infection’. Separate analyses were used to test the effect of sex and mating in females.

| Response Variable | Predictor | Df | F | % Variance Explained | p-value |
| --- | --- | --- | --- | --- | --- |
| Quantitative DCV Shedding | Sex | 1 | 0.67 | 0.28 | 0.42 |
|  | Genetic Background | 9 | 2.52 | 9.48 | 0.009 |
|  | Viral Load | 1 | 5.03 | 4.21 | 0.007 |
|  | DPI | 2 | 0.23 | 0.095 | 0.63 |
|  | Sex*Genetic Background | 9 | 1.73 | 6.53 | 0.082 |
|  | Mating | 1 | 0.22 | 0.098 | 0.64 |
|  | Genetic Background | 8 | 1.44 | 5.82 | 0.17 |
|  | Viral Load | 1 | 11.2 | 10.1 | <0.0001 |
|  | DPI | 2 | 0.18 | 0.08 | 0.67 |
|  | Mating*Genetic Background | 8 | 2.46 | 8.87 | 0.014 |

### Variation in Transmission Potential, *V*

We incorporated the lifespan and virus shedding data described above alongside previously gathered data on genetic and sex-specific variation in social aggregation to calculate individual disease transmission potential, *V* (Lloyd-Smith et al., 2005; VanderWaal & Ezenwa, 2016). As a result of many flies not shedding DCV (Figure 3a), the distribution of transmission potential, *V*, was zero-inflated (Figure 4a). Zero values of *V* represent individuals with no transmission risk (Figure 4a), as flies that shed no virus had no transmission potential, irrespective of their aggregation and lifespan. The distribution of *V* was also characterised by a right-extreme tail, comprised of individuals with high-risk transmission potentials relative to the population average (Figure 4a). Qualitative variation in *V* (the proportion of flies where *V*>0) differed between males and females with the extent of this difference also determined by genetic background (Figure 4b; Table 7). Sex (0.28%), genetic background (2.3%) and the interaction between the two (2.83%) explained relatively little deviance in our models (Figure 5; Table 7). In quantitative variation in *V*, sex explained <1%, while genetic background and its interaction with sex explained 4.13% and 11.4% of variance respectively (Figure 5; Table 8).

**Table 7.** Model outputs for the logistic regression analysis conducted on qualitative *V* (the proportion of non-zero *V* values).

| Response Variable | Predictor | Df | $\chi^2$ | % Deviance Explained | p-value |
| --- | --- | --- | --- | --- | --- |
| Qualitative $V$ | Sex | 1 | 4.58 | 0.28 | 0.032 |
|  | Line | 9 | 38.2 | 2.30 | <0.0001 |
|  | Sex*Line | 9 | 47.0 | 2.83 | <0.0001 |

**Table 8.** Model outputs for the GLM analysis conducted on quantitative *V* (the subset with non-zero *V* values).

| Response Variable | Predictor | Df | F | % Variance Explained | p-value |
| --- | --- | --- | --- | --- | --- |
| Quantitative $V$ | Sex | 1 | 0.077 | 0.01 | 0.78 |
|  | Line | 9 | 2.51 | 4.13 | 0.008 |
|  | Sex*Line | 9 | 6.94 | 11.4 | <0.0001 |

## Discussion

We identified genetic and sex-specific variation in three key outcomes of DCV infection: lifespan following infection, virus shedding, and virus load. When combined with social aggregation data, this variation resulted in genetic and sex-specific variation in individual transmission potential, *V*. While all of these outcomes of infection influence transmission potential, due to many individuals not shedding any virus, virus shedding exerted more influence over V than variation in lifespan following infection and social aggregation. Due to this central role, below we discuss potential explanations for the effect of mating, as well as genetic and sex-specific variation on virus shedding, and link these to genetic and sex-specific variation in *V*.

### The effect of host genetic background in generating heterogeneity in transmission

The genetic variation in virus shedding affected both qualitative and quantitative variation in DCV shedding. As the distributions of neither social aggregation nor lifespan following infection were zero-inflated, variation in virus shedding appears to be a key determinant of qualitative and quantitative variation in *V*. Differences between genetic backgrounds in qualitative shedding was a key determinant of variation in V, as there is no risk of pathogen transmission in the absence of shedding. Among individuals that shed DCV, between-individual heterogeneity in *V* was achieved through different routes. Some genetic backgrounds, such as males from RAL-818, showed a high proportion of individuals that are likely to spread DCV (Figure 4b), but only to relatively few individuals (Figure 4c). Conversely, other groups, such as females of the RAL-380 genetic backgrounds, showed one of the lowest proportions of individuals able to achieve transmission (Figure 4b), but the individuals that did achieve transmission include outliers with values of V that were orders of magnitude higher than the population average (Figure 4c).

Quantitative and qualitative variation in DCV shedding differed in how they were affected by host genetic background. Qualitative variation was affected by genetic background as part of an interaction with host sex, while this interaction has no significant effect on quantitative DCV shedding (Tables 5 & 6). Similar differences are seen in the amount of deviance and variance genetic background explains in models of qualitative and quantitative variation in DCV shedding. Genetic background accounts for only 2.27% of deviance in qualitative DCV shedding whereas it accounts for 9.48% of the variance in quantitative DCV shedding (Figure 5). Genetic variation therefore appears to play an important role in determining shedding and affects qualitative and quantitative shedding in different ways. Similar effects of genetic backgrounds on parasite shedding have been reported in the Ramshorn snail species, *Biomphlamaria glabrata*, during infection with *Schistosoma mansoni.* Genetic backgrounds differ in how many parasite eggs they shed and how quickly they start shedding after infection (Tavalire et al., 2016). The differences we see in the proportion of flies shedding DCV may be caused by a similar pattern of variation in individual’s delaying virus shedding. Delaying the onset of shedding could be affected by a range of DCV infection symptoms. These include paralysis of muscles in the crop organ of the foregut, abdominal swelling, broad nutritional stress and reduced defecation rate (Chtarbanova et al., 2014).

Genetic background also appears to play a key role in transmission potential, we detected a significant effect on both qualitative and quantitative variation in *V*. The amount of deviance and variance explained by genetic background does not hugely differ (2.3% and 4.13%, respectively). However, when part of an interaction with sex, genetic background accounts for 11.4% of the variance in quantitative variation in DCV shedding, whereas this same interaction only accounts for 2.83% of the deviance in qualitative variation in shedding (Figure 5). Alongside other studies, this highlights the potential significance of genetic variation in pathogen shedding to generating transmission heterogeneity. For example, genetic variation in transmission was demonstrated using families of turbot fish (*Scophthalmus maximus*) which produced outbreaks that differed in how quickly individuals showed symptoms of infection and died (Anacleto et al., 2019). Shedding may underlie this genetic variation in transmission as it was not directly measured and there were no significant differences in infection duration and contact rate (Anacleto et al., 2019). Common garden experiments have revealed shedding dynamics capable of influencing the population-level transmission dynamics of wild populations of the plant, *Plantago lanceolata*. In controlled laboratory settings, multi-strain coinfection was shown to increase the number of spores released of the fungal pathogen, *Podosphaera plantaginis*. Measures of natural populations have also demonstrated outbreak severity increases at higher levels of coinfection (Susi, Barrès, Vale, & Laine, 2015). The relationship between spore shedding and coinfection has also been shown to be affected by host genotype, with genotypes significantly differing in the number of spores released over a number of days post-infection (Susi, Vale, & Laine, 2015). Genetic variation in transmission potential has also been demonstrated in the freshwater ciliate, *Paramecium caudatum*, following *Holospora undulata* infection (Fellous et al., 2012). The genotype of the first individual to be infected was a key determinant of pathogen transmission as host genotype appears to affect the infectious potential of shed pathogens (Fellous et al., 2012). *H. undulata* infectiousness increases with host population density, as reduced variation in contact rate makes infectiousness the primary determinant of transmission (Magalon et al., 2010).

### The effect of host sex in generating heterogeneity in transmission

We also observed clear qualitative and quantitative differences in *V* between males and females, which is suggestive of sex-specific variation in disease transmission. While the extent of any difference between males and females is also determined by genetic background, a greater proportion of males tend to transmit DCV than females across these backgrounds. In DCV shedding, a greater proportion of males from several genetic backgrounds (RAL-379, RAL-738 and RAL-818) shed DCV than females (Figure 3a). Interestingly, we see significant sex-specific differences in qualitative, but not quantitative, variation in DCV shedding. Other work has also shown a number of sex differences in pathogen and parasite shedding (Sanchez, Devevey, & Bize, 2011; Sheridan, Poulin, Ward, & Zuk, 2000; Thompson, Gipson, & Hall, 2017). Often these biases link to differences in the selection pressures applied by sexual reproduction (Duneau & Ebert, 2012). Comparisons of mated and virgin female flies revealed mating effects which produced quantitative, but not qualitative, differences in shedding. While we did not measure *V* in virgin females, this mating effect may offer explanations for the sex differences seen in shedding and therefore *V*.

Sex-specific variation in qualitative differences in shedding exerts a significant influence over shedding (Figure 3a). It is important to note however, that in isolation, sex accounts for a miniscule 0.64% of the deviance in qualitative variation in shedding. Sex appears to play a more important role in conjunction with genetic background, the interaction between the two explaining 3.07% of deviance (Figure 5). While significant, sex-specific variation may play a relatively minor role in shedding. A variety of factors appear to underlie sex-differences in shedding across host-pathogen systems. For example, male-biased infection is common to many mammal hosts but generally absent from arthropod hosts (Sheridan et al., 2000). In the water flea, *Daphnia magna*, parasite spores are released into the environment upon death and females have been shown to release significantly more than males (Thompson et al., 2017). In the vole, *Microtus gryalis*, the faeces of females contains significantly more parasite eggs than that of males (Sanchez et al., 2011). Given that we see female-biased mortality to DCV infection (Figure 1a), it is perhaps surprising that shedding is not also female-biased. This could be due to shedding being measured during the first three days of infection, whereas mortality occurred much later. We might therefore see sex-differences in shedding during the later stages of infection.

Both the qualitative and quantitative differences in *V* between males and females were determined alongside genetic background. While sex explained very little deviance and variance in qualitative and quantitative variation in *V* (Figure 6), its interaction with genetic background explained 2.83% and 11.4 %, respectively. Sex could therefore be an important source of variation in individual disease transmission. Sex differences in transmission or virus shedding, lifespan and social aggregation are commonly observed in a wide range of species (Duneau & Ebert, 2012; Ferrari, Cattadori, Nespereira, Rizzoli, & Hudson, 2004; Kaltz & Shykoff, 2001; Sanchez et al., 2011). Sex-specific variation has been relatively well-studied because sexes are easily distinguished in the wild, and examples of sexual dimorphism in physiological and behavioural traits are relatively common (Duneau and Ebert, 2012). Many mammalian hosts exhibit male-biased transmission (Ezenwa et al., 2016; Grear, Luong, & Hudson, 2012; Luong, Grear, & Hudson, 2009; Rhines, 2013), although there are exceptions of female-bias (Sanchez et al., 2011). In the white-footed mouse, *Peromyscus leucopus*, male-biased transmission is thought to be driven by sex differences in contact network connectivity, which has been linked to testosterone production (Foo, Nakagawa, Rhodes, & Simmons, 2017; Grear et al., 2012). Testosterone may be particularly relevant to transmission as its immunosuppressive (Foo et al., 2017) effects may also alter infectiousness and infection duration.

### Female Mating Status in Shedding

Mated and virgin females did not qualitatively differ in DCV shedding; however, individuals did exhibit quantitative variation in shedding. While only 0.098% of the variance in quantitative shedding was explained by mating, the interaction between mating and genetic background explained 8.87% of the variance (Figure 5). This suggests that alongside host genetic background, mating might exert an important level of influence over shedding. One potential explanation for this mating effect are post-mating physiological changes in the intestine that can increase in defecation rates (Apger-McGlaughon and Wolfner, 2013). However, if this change is responsible for the significant effect of female mating, the virgin females from particular genetic backgrounds that shed more than mated females (Figure 3b) may be tolerant to these physiological changes. Relatively few have considered how mating affects aspects of disease transmission outside of contact rates (Altizer et al., 2003; Thrall et al., 2000). Particularly alongside other work in *Drosophila* that has demonstrated female-specific costs of infection (Kubiak and Tinsley, 2017; Short et al., 2012), this result highlights the importance of mating-induced physiological changes to transmission heterogeneity.

The difference between qualitative and quantitative variation in shedding relates to assumptions we make regarding how often DCV is shed. If DCV is always present in shedding, measures of zero reflect quantities of virus that are below the detection threshold of qPCR. While this could result in infectious individuals evading detection, as oral infection typically requires very high dosage (Gupta et al., 2017; Palmer et al., 2018), low-titre zero-values pose a smaller transmission risk. If DCV is not always shed, within-individual variation in when shedding occurs could be central to transmission heterogeneity (Chen, Sanderson, & Lanzas, 2013). This is particularly relevant to our study as virus shedding was only measured at a single time point per fly, and we do not know how shedding, and therefore *V*, may change over time. Within-host, temporal variation in shedding is observed in a range of host-pathogen systems (Chen et al., 2013; Matthews et al., 2006; Mideo, Alizon, & Day, 2008). For example, avian hosts tend to shed more parasites during the late afternoon (Brawner III and Hill, 1999; Martinaud et al., 2009).

By combining measures of virus shedding, lifespan and social aggregation into a simple framework our work demonstrates that genetic and sex-specific variation can affect individual heterogeneity in disease transmission potential. We also show that genetic and sex-specific variation, as well as mating, can produce variation outcomes of infection. Alongside its interaction with sex, genetic background explains 5.41% of qualitative, and 15.54% of quantitative, individual variation in transmission potential. While our results do not implicate a particular genetic background, males generally present a greater transmission risk than females. In addition to highlighting high-risk individuals, our results are congruous with the observation that the majority of infected individuals produce very few, if any, secondary cases of infection. Non-infectious individuals are particularly relevant to predicting outbreaks of infectious disease as they obscure high-risk individuals in traditional, population-wide estimations of outbreak risk. Our findings demonstrate the benefit of using a model laboratory system as well established as *D. melanogaster* to study disease transmission. The number of available protocols and methodologies are central to considering multiple traits central to disease transmission and holistically studying their underlying determinants.

## Acknowledgements

J.A.S-J was funded by a NERC E3 DTP PhD studentship awarded to the University of Edinburgh. P.F.V was supported by a Branco Weiss fellowship (https://brancoweissfellowship.org/) and a Chancellor’s Fellowship (School of Biological Sciences, University of Edinburgh).

We would like to thank F. Waldron for assistance with RNA extraction and troubleshooting as well as V. Gupta, K. Monteith, H. Borthwick, H. Cowan, and A. Reid for technical assistance and media preparation.

## Supplementary Figures and Tables

**Figure S1.**
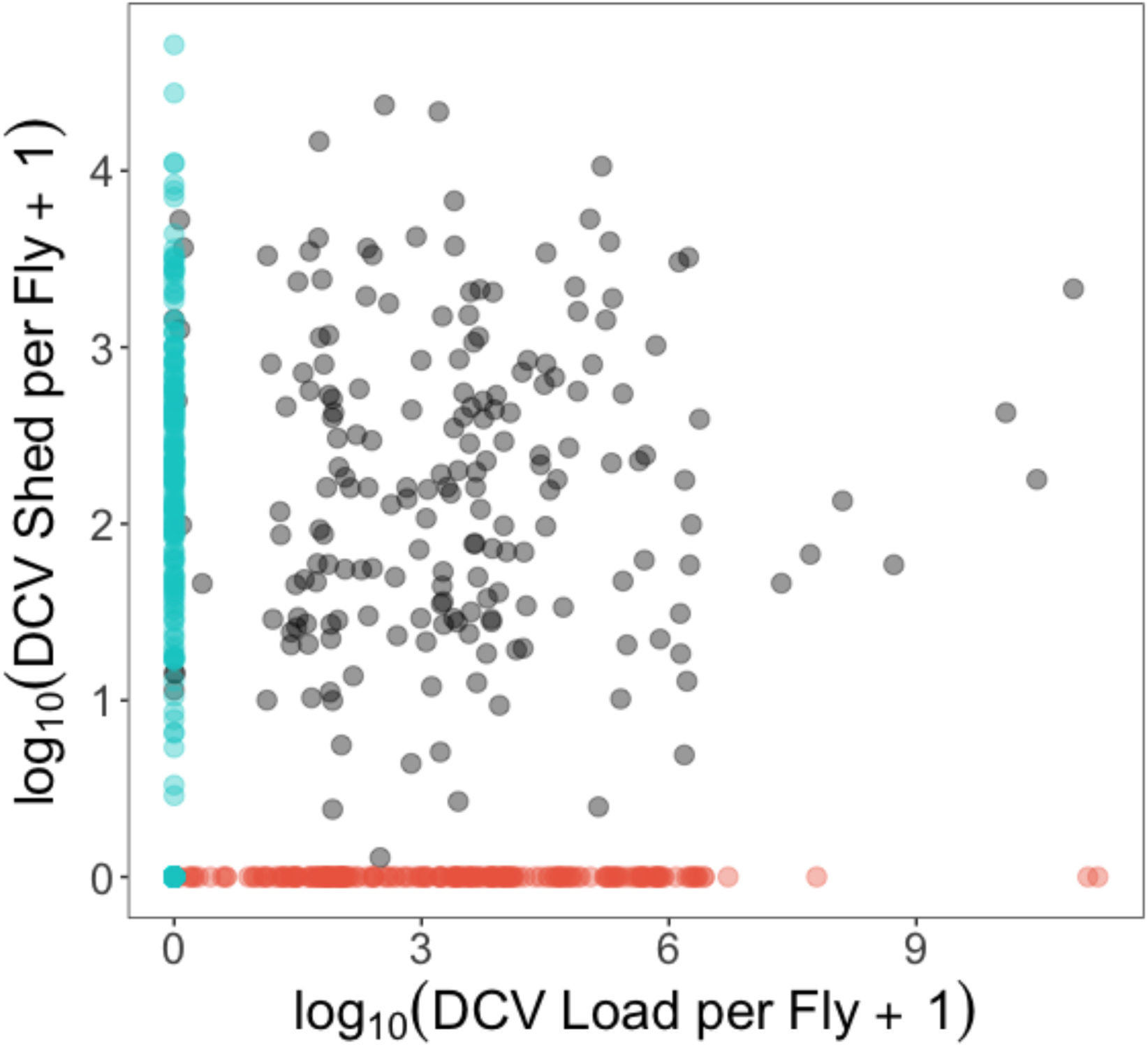
The relationship between the viral load of flies and the amount of virus they shed into their environment. The two distinct phenotypes, where individuals show a zero-value for shedding or load and a positive-value for the other trait, are marked by blue (supersponges) or red (supershedders).

**Table S1.** The number of flies measured for lifespan and viral load at death for each combination of genetic background and sex/female mating status.

|  | 59 | 75 | 138 | 373 | 379 | 380 | 502 | 738 | 765 | 818 |
| --- | --- | --- | --- | --- | --- | --- | --- | --- | --- | --- |
| Male | 20 | 20 | 18 | 19 | 17 | 18 | 17 | 19 | 18 | 19 |
| Female | 17 | 20 | 13 | 19 | 19 | 20 | 19 | 18 | 15 | 18 |
| Virgin Female | 20 | 18 | 7 | 18 | 20 | 19 | 19 | 20 | 16 | 20 |

**Table S2.** The number of viral load samples for each treatment group (a) 1 DPI, (b) 2 DPI and (c) 3 DPI.

|  |  |  |  |  |  |  |  |  |  |  |  |
| --- | --- | --- | --- | --- | --- | --- | --- | --- | --- | --- | --- |
| (a) |  | 59 | 75 | 138 | 373 | 379 | 380 | 502 | 738 | 765 | 818 |
|  | Male | 12 | 11 | 11 | 12 | 11 | 11 | 12 | 11 | 11 | 6 |
|  | Mated Female | 13 | 12 | 11 | 12 | 12 | 12 | 11 | 12 | 12 | 12 |
|  | Virgin Female | 12 | 14 | NA | 15 | 12 | 13 | 12 | 12 | 7 | 12 |
| (b) |  | 59 | 75 | 138 | 373 | 379 | 380 | 502 | 738 | 765 | 818 |
|  | Male | 12 | 12 | 11 | 12 | 11 | 12 | 11 | 12 | 12 | 6 |
|  | Mated Female | 14 | 12 | 11 | 11 | 12 | 11 | 11 | 11 | 12 | 13 |
|  | Virgin Female | 12 | 14 | NA | 14 | 12 | 11 | 13 | 10 | 9 | 12 |
| (c) |  | 59 | 75 | 138 | 373 | 379 | 380 | 502 | 738 | 765 | 818 |
|  | Male | 11 | 12 | 12 | 12 | 12 | 11 | 12 | 10 | 12 | 7 |
|  | Mated Female | 11 | 11 | 10 | 13 | 12 | 11 | 13 | 11 | 12 | 12 |
|  | Virgin Female | 11 | 13 | NA | 13 | 12 | 13 | 11 | 12 | 8 | 11 |

**Table S3.** The number of non-zero viral load samples for each treatment group (a) 1 DPI, (b) 2 DPI and (c) 3 DPI.

|  |  |  |  |  |  |  |  |  |  |  |  |
| --- | --- | --- | --- | --- | --- | --- | --- | --- | --- | --- | --- |
| (a) |  | 59 | 75 | 138 | 373 | 379 | 380 | 502 | 738 | 765 | 818 |
|  | Male | 5 | 9 | 2 | 5 | 4 | 6 | 8 | 6 | 5 | 3 |
|  | Mated Female | 6 | 5 | 5 | 10 | 8 | 9 | 2 | 9 | 3 | 9 |
|  | Virgin Female | 4 | 11 | NA | 11 | 7 | 9 | 4 | 3 | 6 | 4 |
| (b) |  | 59 | 75 | 138 | 373 | 379 | 380 | 502 | 738 | 765 | 818 |
|  | Male | 1 | 5 | 3 | 1 | 1 | 3 | 7 | 6 | 1 | 4 |
|  | Mated Female | 8 | 4 | 5 | 5 | 2 | 1 | 1 | 3 | 4 | 1 |
|  | Virgin Female | 1 | 3 | NA | 9 | 7 | 7 | 6 | 1 | 5 | 4 |
| (c) |  | 59 | 75 | 138 | 373 | 379 | 380 | 502 | 738 | 765 | 818 |
|  | Male | 1 | 5 | 5 | 5 | 7 | 7 | 1 | 4 | 7 | 6 |
|  | Mated Female | 5 | 4 | 7 | 4 | 5 | 4 | 1 | 7 | 9 | 2 |
|  | Virgin Female | 1 | 1 | NA | 6 | 6 | 6 | 5 | 6 | 3 | 3 |

**Table S4.** The number of virus shedding samples for each treatment group (a) 1 DPI, (b) 2 DPI and (c) 3 DPI.

|  |  |  |  |  |  |  |  |  |  |  |  |
| --- | --- | --- | --- | --- | --- | --- | --- | --- | --- | --- | --- |
| (a) |  | 59 | 75 | 138 | 373 | 379 | 380 | 502 | 738 | 765 | 818 |
|  | Male | 10 | 11 | 10 | 12 | 11 | 10 | 12 | 10 | 10 | 5 |
|  | Mated Female | 13 | 12 | 11 | 11 | 12 | 12 | 11 | 12 | 12 | 12 |
|  | Virgin Female | 11 | 14 | NA | 15 | 12 | 13 | 12 | 12 | 7 | 12 |
| (b) |  | 59 | 75 | 138 | 373 | 379 | 380 | 502 | 738 | 765 | 818 |
|  | Male | 12 | 12 | 11 | 11 | 10 | 12 | 11 | 12 | 12 | 7 |
|  | Mated Female | 14 | 12 | 11 | 11 | 12 | 11 | 11 | 9 | 12 | 13 |
|  | Virgin Female | 11 | 13 | NA | 13 | 12 | 11 | 13 | 11 | 9 | 12 |
| (c) |  | 59 | 75 | 138 | 373 | 379 | 380 | 502 | 738 | 765 | 818 |
|  | Male | 10 | 11 | 12 | 12 | 11 | 11 | 12 | 10 | 9 | 7 |
|  | Mated Female | 11 | 9 | 10 | 13 | 9 | 9 | 13 | 8 | 12 | 11 |
|  | Virgin Female | 10 | 12 | NA | 13 | 12 | 12 | 7 | 11 | 7 | 10 |

**Table S5.** The number of non-zero virus shedding samples for each treatment group (a) 1 DPI, (b) 2 DPI and (c) 3 DPI.

|  |  |  |  |  |  |  |  |  |  |  |  |
| --- | --- | --- | --- | --- | --- | --- | --- | --- | --- | --- | --- |
| (a) |  | 59 | 75 | 138 | 373 | 379 | 380 | 502 | 738 | 765 | 818 |
|  | Male | 0 | 7 | 4 | 6 | 3 | 7 | 4 | 6 | 2 | 4 |
|  | Mated Female | 5 | 4 | 3 | 6 | 6 | 5 | 1 | 6 | 7 | 4 |
|  | Virgin Female | 4 | 6 | NA | 8 | 6 | 4 | 6 | 3 | 5 | 5 |
| (b) |  | 59 | 75 | 138 | 373 | 379 | 380 | 502 | 738 | 765 | 818 |
|  | Male | 10 | 7 | 6 | 5 | 9 | 9 | 5 | 11 | 7 | 7 |
|  | Mated Female | 8 | 4 | 9 | 7 | 8 | 3 | 6 | 3 | 4 | 4 |
|  | Virgin Female | 5 | 6 | NA | 7 | 10 | 5 | 8 | 8 | 6 | 6 |
| (c) |  | 59 | 75 | 138 | 373 | 379 | 380 | 502 | 738 | 765 | 818 |
|  | Male | 4 | 4 | 8 | 3 | 9 | 2 | 7 | 2 | 3 | 6 |
|  | Mated Female | 7 | 3 | 6 | 7 | 2 | 2 | 3 | 3 | 4 | 5 |
|  | Virgin Female | 4 | 4 | NA | 7 | 7 | 4 | 4 | 5 | 2 | 4 |

**Table S6.** Summaries of the logistic regression and GLMs used to analyse the response variables of our experiments. All interactions are fully-factorial and marked using an asterisk (*).

| Response Variable | Analysis | Predictors |
| --- | --- | --- |
| Lifespan | GLM | Sex * Genetic Background * VLAD<br>Mating * Genetic Background * VLAD |
| VLAD | GLM | Sex * Genetic Background<br>Mating * Genetic Background |
| Qualitative Load | Logistic Regression | Sex * Genetic Background + DPI<br>Mating * Genetic Background + DPI |
| Quantitative Load | GLM | Sex * Genetic Background + DPI<br>Mating * Genetic Background + DPI |
| Qualitative Shed | Logistic Regression | Sex * Genetic Background + Quant. Load + DPI<br>Mating * Genetic Background + Quant. Load + DPI |
| Quantitative Shed | GLM | Sex * Genetic Background + Quant. Load + DPI<br>Mating * Genetic Background + Quant. Load + DPI |
| Qualitative V | Logistic Regression | Sex * Genetic Background |
| Quantitative V | GLM | Sex * Genetic Background |

